# *TERT* expression is associated with metastasis from thin primaries, exhausted CD4+ T cells in melanoma and with DNA repair across cancer entities

**DOI:** 10.1101/2023.01.25.525565

**Authors:** Christina Katharina Kuhn, Jaroslawna Meister, Sophia Kreft, Mathias Stiller, Sven-Holger Puppel, Anne Zaremba, Björn Scheffler, Vivien Ullrich, Torsten Schöneberg, Dirk Schadendorf, Susanne Horn

**Affiliations:** Rudolf Schönheimer Institute of Biochemistry, University of Leipzig, Medical Faculty, Johannisallee 30, 04103 Leipzig, Germany; Institute for Clinical Diabetology, German Diabetes Centre, Leibniz Centre for Diabetes Research at Heinrich Heine University Düsseldorf, 40225 Düsseldorf, Germany; Department of Dermatology, University Hospital Essen, University Duisburg-Essen, and German Cancer Consortium (DKTK) partner site Essen/Düsseldorf, 45147 Essen, Germany; Institute of Pathology, University of Leipzig Medical Center, Liebigstraße 26, 04103 Leipzig, Germany; DKFZ-Division Translational Neurooncology at the West German Cancer Center (WTZ), University Hospital Essen/University of Duisburg-Essen, 45147 Essen, Germany

**Keywords:** telomerase, transcriptomics, survival, immune checkpoint inhibition, metastasis

## Abstract

Telomerase reverse transcriptase (*TERT*) promoter mutations occur frequently in cancer, have been associated with increased *TERT* expression and cell proliferation, and could potentially influence therapeutic regimens for melanoma. As the role of *TERT* expression in malignant melanoma and the non-canonical functions of TERT remain understudied, we aimed to extend the current knowledge on both types of *TERT* alterations with respect to survival, further clinical and molecular parameters. Using multivariate models, *TERT* alterations were not consistently associated with survival in melanoma cohorts under immune checkpoint inhibition. The presence of CD4+ T cells increased with *TERT* expression and correlated with the expression of exhaustion markers. While the frequency of promoter mutations did not change with Breslow thickness, *TERT* expression was increased in metastases arising from thinner primaries. Enrichment analyses of single-cell RNA-seq showed *TERT* expression is associated with genes involved in cell migration and dynamics of the extracellular matrix, supporting the role of *TERT* during invasion and metastasis. Co-regulated genes in several bulk tumors and single-cell RNA-seq cohorts also indicated non-canonical functions of *TERT* related to mitochondrial DNA stability and nuclear DNA repair in line with increased *TERT* expression during chromothripsis (PCAWG cohort) and under hypoxic conditions (PCAWG and SKCM cohorts). Also in glioblastoma (Klughammer and PCAWG cohorts), *TERT* was co-expressed with DNA repair genes. Our results thus indicate a relevance of *TERT* expression in melanoma metastasis, T cell dysfunction and DNA repair across cancer entities.

**Significance:** In addition to the frequently occurring *TERT* promoter mutations, we test *TERT* expression with respect to clinical and molecular associates, extending the canonical role of *TERT* in melanoma and other cancer entities.

## Introduction

Telomere maintenance mechanisms play a critical role in cellular survival as they are known to enhance the proliferative capacity of cancer cells [1]. Frequent mutations in the promoter region of the telomerase reverse transcriptase (*TERT*) gene were shown to be associated with the upregulation of telomerase expression [2–4]. *TERT* promoter mutations have been proposed as independent prognostic markers in non-acral cutaneous melanoma, since they correlated in a multivariate analysis with decreased overall survival (OS) without analysis of therapy [5]. The co-occurrence of *BRAF* or *NRAS* mutations with *TERT* promoter mutations was also prognostic for poor disease free survival (DFS) in multivariate analyses in primary melanoma [6]. *TERT* promoter mutations may serve as predictive biomarkers as they were independently associated with improved survival in patients receiving BRAF/MEK inhibitor therapy in univariate and multivariate analyses [7,8]. Also, patients undergoing anti-CTLA4 therapy showed better overall survival with *TERT* promoter mutations, implying its potential as a predictive marker for immune checkpoint inhibitor (ICI) therapy [9]. Also, *TERT* expression has been suggested to be an independent prognostic marker associated with poor survival in human cancer patients beyond melanoma [10–12]. However, many variables, such as mutational load, tumor heterogeneity, and prior therapies, could be appended to existing multivariate models to test additional potentially stronger predictors.

The extent to which genetic alterations of *TERT* play a role in cancer progression and the molecular mechanisms affected by reactivation of *TERT* transcription vary widely among cancer entities, making it crucial to understand the underlying mechanisms [13]. One way cancer cells regain their proliferative ability is by forming a new ETS binding motif at the promoter mutation [2]. This binding motif attracts transcription factors, such as TCF1, Myc, Wnt/β-catenin, NF-kB, and GAPB, which are responsible for regulating numerous cellular processes including tumorigenesis [2,14]. The canonical function of *TERT* is telomere maintenance, however, non-canonical functions of *TERT* have been discussed, e.g. related to reactive oxygen species (ROS), mitochondria and aging. TERT is imported into the mitochondria, where it binds mitochondrial DNA, protecting it from oxidative stress–induced damage, reducing apoptosis in human endothelial cells [15]. Furthermore, besides the full-length expressed *TERT,* alternatively spliced isoforms exist, which lack telomerase activity and may account for up to 10% of endogenous *TERT*. These are continuously expressed after telomerase silencing during human embryonic development [16] and are potentially associated with telomere-independent effects [17].

Here, we test if *TERT* promoter mutations and *TERT* expression are associated with OS and progression free survival (PFS) under ICI, as well as clinicopathological and demographic parameters in multiple highly annotated melanoma cohorts with exome- and transcriptome sequencing. To examine the non-canonical functions of *TERT*, we investigate associations with ROS-related genes, hypoxia, chromothripsis, and *TERT* isoform expression. Using molecular pathway analyses, we replicate known and identify new signaling pathways regulated with genetically modified *TERT* in bulk tumors, cell lines, and single-cells. Tumor immune signatures are estimated to elucidate the role of *TERT* during tumor rejection. Finally, our analyses are extended to additional cancer entities from the Pan-Cancer Analysis of Whole Genomes (PCAWG) cohort and glioblastoma to determine the signaling pathways co-regulated with *TERT* across entities.

## Materials and Methods

### NGS data analysis

We analyzed several bulk RNA-seq cohorts with respective survival data (Table 1, Supplementary Methods). These included five melanoma cohorts with samples taken prior to or early-on ICI therapy (Gide, Hugo, Liu, Riaz, Van Allen cohorts), one melanoma cohort without ICI therapy (SKCM), two cross-entity datasets: cell lines from the Cancer Cell Line Encyclopedia (CCLE), and bulk tumors of the Pan-Cancer Analysis of Whole Genomes (PCAWG). We further included melanoma single-cell RNA-seq (scRNA-Seq) and glioblastoma tumors in our workflow (Supplementary Fig. S1). In brief, we extended the available gene expression levels with additional metadata, as described below and in the supplementary methods. We annotated published *TERT* promoter mutations in the RNA-seq datasets of two melanoma cohorts (Van Allen, SKCM) and two cross-entity datasets (CCLE and PCAWG). We added immune signatures for the six melanoma bulk RNA-seq cohorts using TIMER 2.0 [18]). Isoform analysis was performed for the Van Allen, SKCM and PCAWG cohorts. To analyze mitochondria-related features in the PCAWG and SKCM cohorts, we further integrated measurements for hypoxia, chromothripsis [19], and mitochondrial mutations [19,20]. Hypoxia scores for PCAWG were determined as in Buffa et. al 2010 [21], and hypoxia scores for the SKCM cohort were obtained from the cBioPortal (www.cbioportal.org/study/clinicalData?id=skcm_tcga_pan_can_atlas_2018). We reanalyzed the MSK IMPACT Clinical Sequencing Cohort (dataset “MSKCC, Nat Med 2017”) [9] as obtained from www.cbioportal.org annotating the corresponding therapy data from Samstein [22].

**Table 1.**
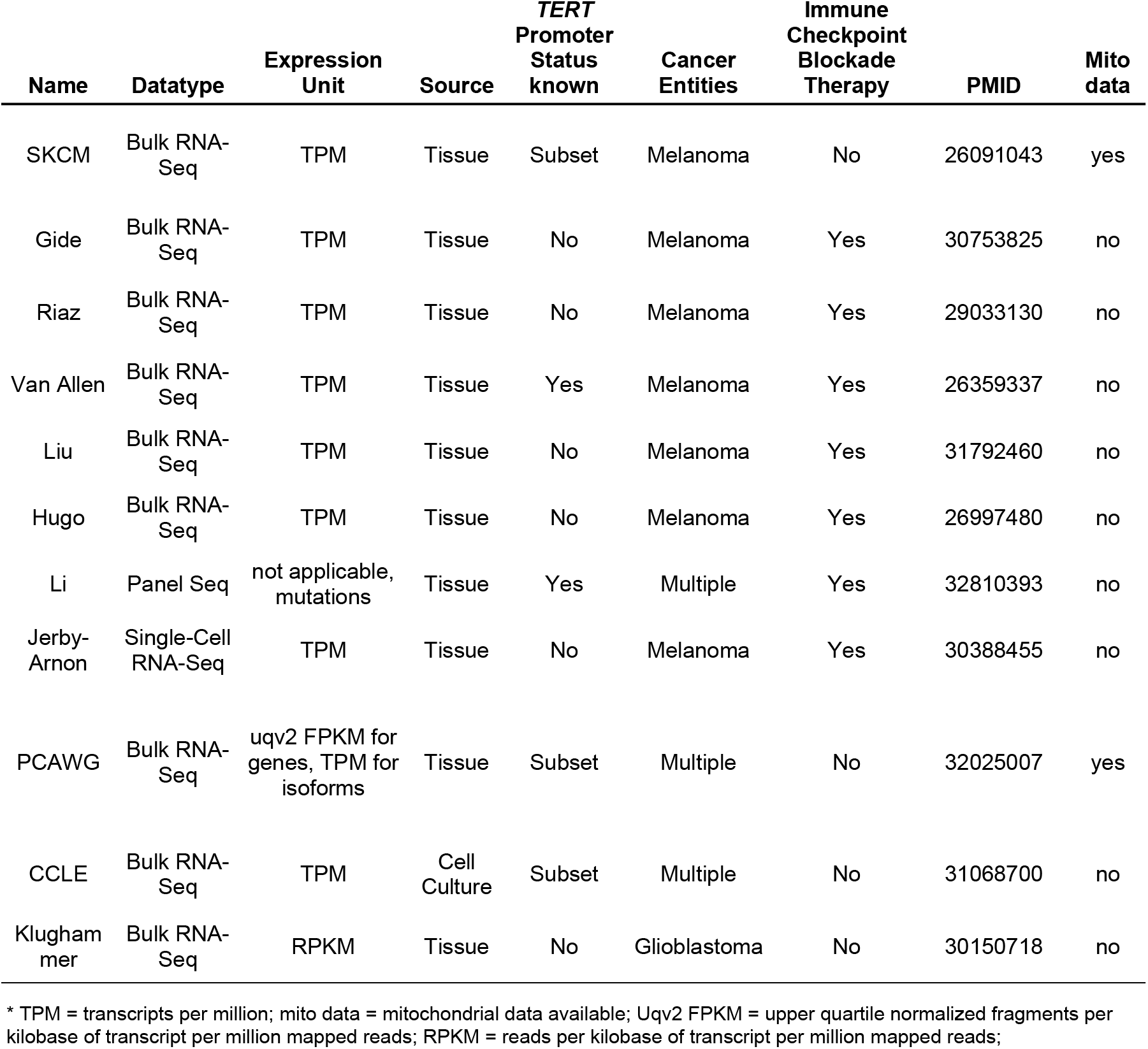
Analyzed cohorts.

### Enrichment analysis

Enrichment analysis of top genes associated with *TERT* promoter mutation and *TERT* expression using bulk RNA-seq cohorts was conducted using the gene set enrichment analysis tool https://maayanlab.cloud/Enrichr/ [23] with default settings. For enrichment analysis in the melanoma single-cell context, the top 100 genes associated with *TERT* expression in the scRNA-seq dataset were queried with Enrichr using default settings. *TERT* promoter mutation status was not available for single-cell data. The top ten results of each enrichment set from gene ontology (GO) pathways, ontologies and cell types were then used for discussion of the most frequently occurring terms.

### *TERT* isoforms analysis

Short names, coding information, biotype, and transcript lengths of *TERT* isoforms were collected from Ensembl (GRCh38.p13) using ENSG gene identifiers (Supplementary Methods). The presence of the mitochondrial targeting protein sequence ‘MPRAPRCRAVRSLLRSHYRE’ [24] in the translated isoforms was analyzed using the UCSC Human Genome Browser. In detail, we verified whether transcripts included the targeting sequence by aligning the protein sequence with the genome (GRCh38.p13) and comparing the start/stop positions with the isoform exons. The isoform expressed in normal control tissues was assessed using the GTEx portal (https://gtexportal.org/home/gene/TERT#gene-transcript-browser-block, Supplementary Fig. S7). We then summed over the measured transcripts per million (TPM) of all isoforms and calculated the proportion of isoforms with mitochondrial targeting sequence. Data on isoforms lacking the targeting sequence were available only for the CCLE and Van Allen cohorts.

### Statistics

Correlation of *TERT* mRNA expression, isoform expression, and clinical variables was tested using Spearman correlations. For comparison between samples with or without *TERT* promoter mutation, two-sided Wilcoxon rank tests were applied. The correlates associated with *TERT* expression or *TERT* promoter mutations were ranked by p-value with FDR control, and overlaps of the gene lists from several cohorts were performed. For survival analysis, the cohorts were divided into low vs. high *TERT* expression groups, with approximately one-third of the patients showing high *TERT* expression. If available, cohorts (SKCM and Van Allen cohort) were also divided into two groups by *TERT* promoter mutation absence/presence. Kaplan-Meier survival curves, log-rank tests, multivariate Cox Proportional-Hazards analyses, and odds ratios were computed using R v.4.0.1 [25].

### Data availability statement

The data used in this study are available within this manuscript and its supplementary files. Previously published, processed NGS data can be accessed at https://tools.hornlab.org/cru337phenotime/.

## Results

### Association of *TERT* promoter mutation status and *TERT* expression with survival

Starting from the genomic level, we analyze the role of *TERT* promoter mutations in malignant melanoma in each of the following sections and extend the analysis to the transcriptomic level by examining the impact of *TERT* expression on clinical and molecular associations. In our univariate and multivariate survival analyses of six bulk RNA-seq cohorts of metastatic melanoma treated with ICI (Gide, Riaz, Van Allen, Liu, Hugo cohorts) and without targeted therapy or ICI (SKCM), *TERT* promoter mutations and *TERT* expression were not associated with survival (Supplementary Fig. S2-4, Supplementary Table S1). PFS under anti-PD1 immunotherapy with pembrolizumab appeared decreased with higher *TERT* expression in only one analyzed cohort with small sample size (n=8 vs. 24, Gide cohort, PFS log rank p=0.038), however, it was not significantly associated in multivariate Cox PH including age, sex and *BRAF* mutation status (p=0.071, Supplementary Table S1A).

In contrast to our finding, an association of *TERT* promoter mutations with survival under anti-CTLA4 therapy was described previously [9], however, no multivariate model was attempted. Therefore, we reanalyzed the melanoma subset of the MSK Impact 2017 cohort [9] and the corresponding therapy data and included tumor mutational burden (TMB), sex, and *BRAFV*600 variants in a multivariate model. The previously described association of *TERT* promoter mutations with survival still holds, albeit with reduced significance (p=0.015 compared to the previous univariate analysis p=0.0007, Supplementary Fig. S5A-C).

### *TERT* expression and increased intratumoral CD4+ T helper 1 cells

The top ten significant immune signatures correlated with *TERT* expression in the SKCM cohort included CD4+ and CD8+ T helper type 1/2 cells, macrophages, neutrophils, and natural killer cells (Supplementary Table S2). None of these signatures were consistently regulated in the same direction of effect in the other five bulk RNA-seq cohorts. However, the CD4+ T helper type 1 cell signature repeatedly increased with *TERT* in five of the six cohorts analyzed here (Fig. 1A). Within a patient population with high type 1 CD4+ T cell estimates, survival did not differ with elevated *TERT* expression (Supplementary Fig. S6).

**Figure 1.**
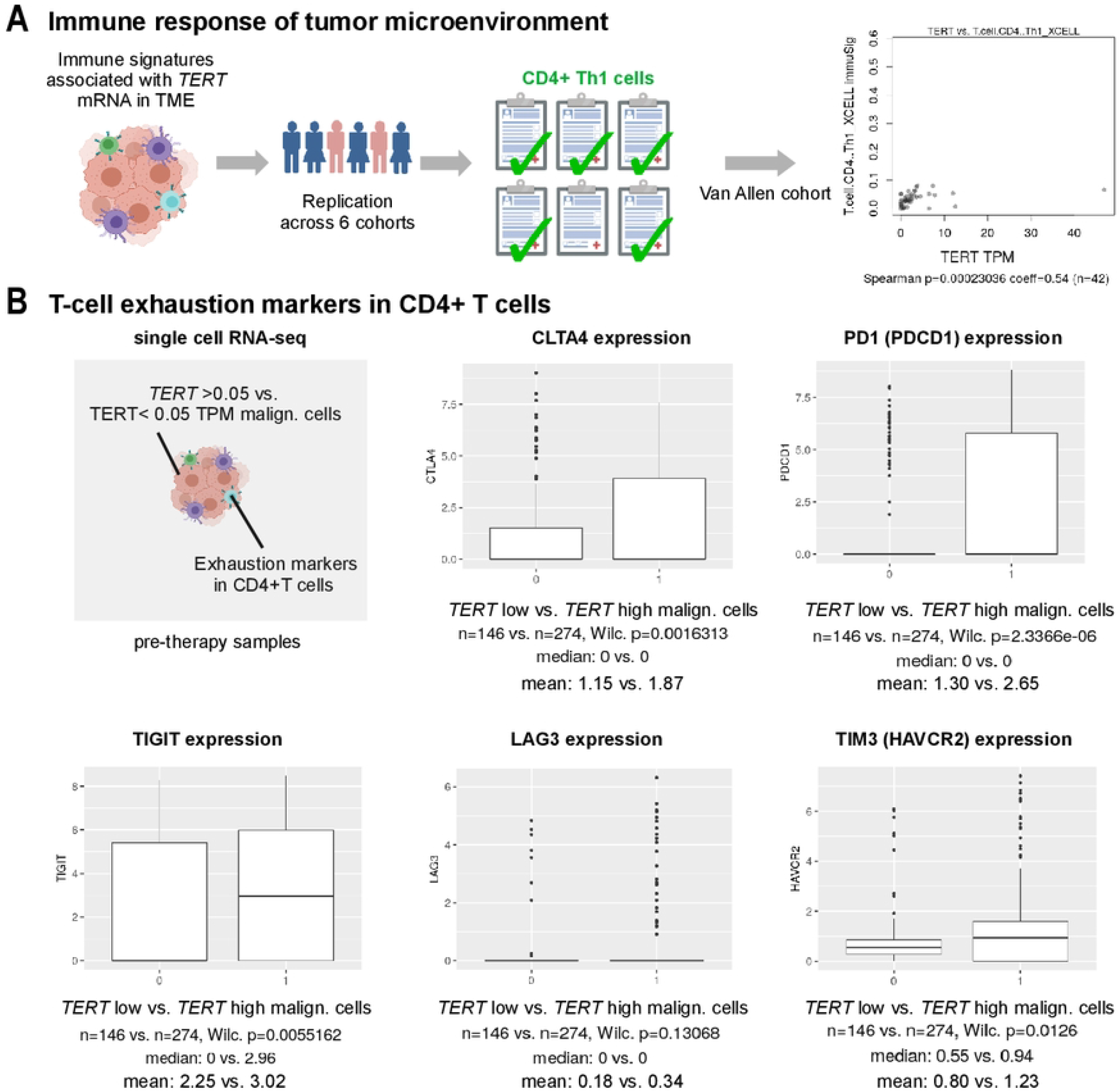
**A,** *TERT* expression and immune signatures in six bulk tumor cohorts. Five out of six cohorts showed correlation of *TERT* expression and CD4+ Th1 T cell immune signature (xCell assay). **B**, Expression of T cell exhaustion markers in single CD4+ T cells of patients with low vs. high *TERT* expression in malignant tumor cells (</>0.05 TPM). Jerby-Arnon cohort, pre therapy samples. Th1: T-helper type 1; TME: Tumor microenvironment; TPM: Transcripts per million; xCell: xCell cell types enrichment analysis.

### Primary tumor thickness

Our examination of clinical variables in association with *TERT* promoter mutations and expression was based on the two largest melanoma cohorts, the SKCM and Liu cohorts (Supplementary Tables S3A-C). There was no obvious difference in the primary tumor thickness between metastatic samples with vs. without *TERT* promoter mutations (Wilcoxon p=0.508, Fig. 2A). Notably, *TERT* expression was higher in metastases of tumors arising from thinner primaries (Spearman p=0.01, coeff=−0.16, Fig. 2A; Wilcoxon p=0.011, Breslow >2 mm: 8.95 TPM vs. Breslow <2 mm: 11.89 TPM, Fig. 2B).

**Figure 2.**
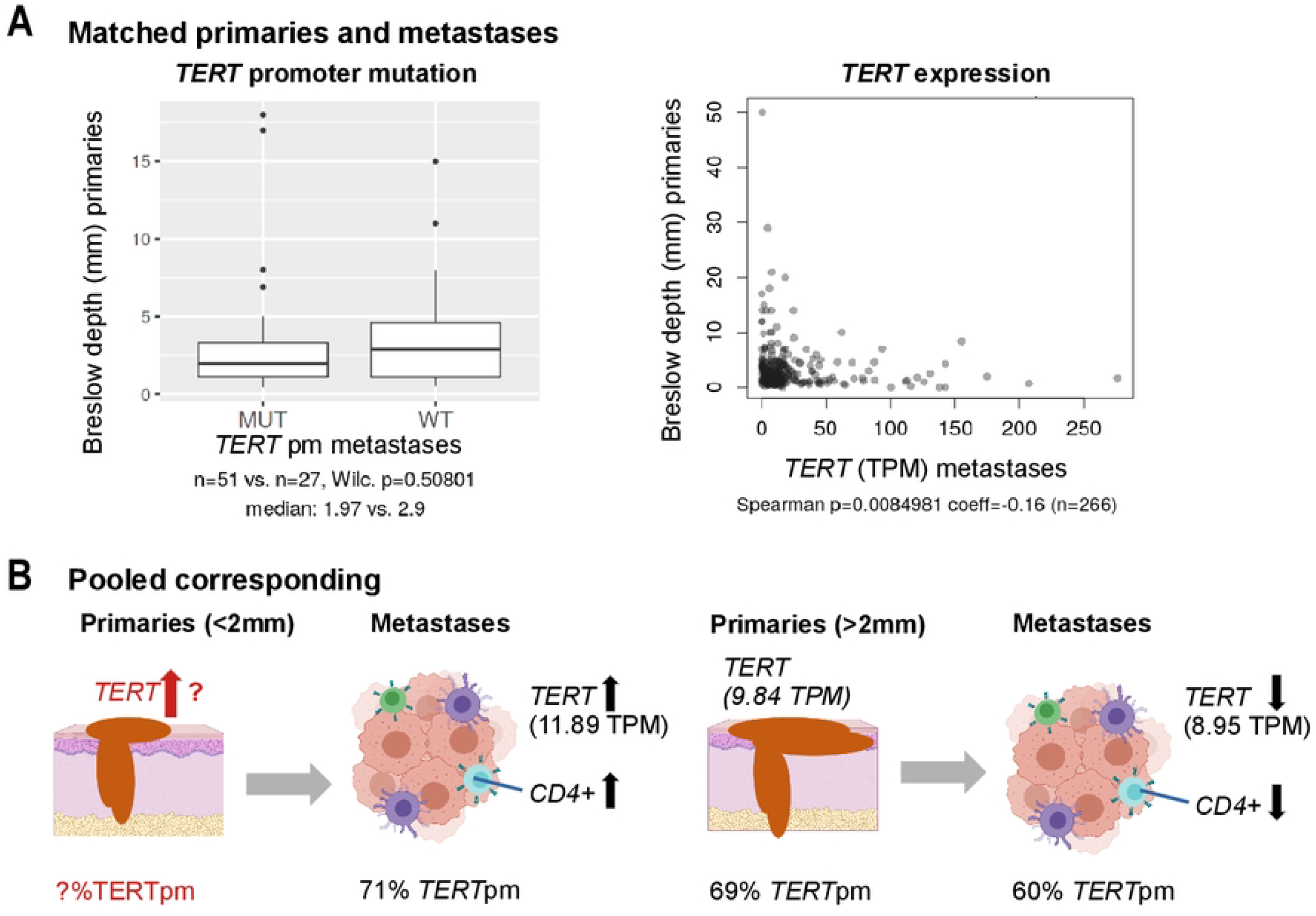
**A,** Breslow depth of primaries with *TERT* promoter mutations and *TERT* expression measured in corresponding metastatic tumors of the SKCM cohort (without one outlier; *TERT* expression >600, for data including outliers see Supplementary Fig. S9; also see Supplementary Table S2D). **B,** *TERT* expression in metastatic tumors arising from thin (Breslow depth <2 mm) and thick tumors (>2 mm). Information on *TERT* alterations in thin primaries not available. *TERT*pm: *TERT* promoter mutations; TPM: Transcripts per million.

### Mutational load

A common pattern of *TERT* promoter mutations and *TERT* expression was an association with more nuclear mutational load. In the entire SKCM cohort, samples with *TERT* promoter mutations showed higher overall mutation counts (Wilcoxon p=0.014, Fig. 3B, not significant for the metastatic subgroup) and higher *TERT* expression was also associated with mutation counts (Spearman p=0.002, coeff=0.16, Fig. 3B, metastatic subgroup: Spearman p=0.017, coeff=0.14), however, not in the Liu cohort (Spearman: 0.194, coeff=0.12, Supplementary Table S3A-B).

**Figure 3.**
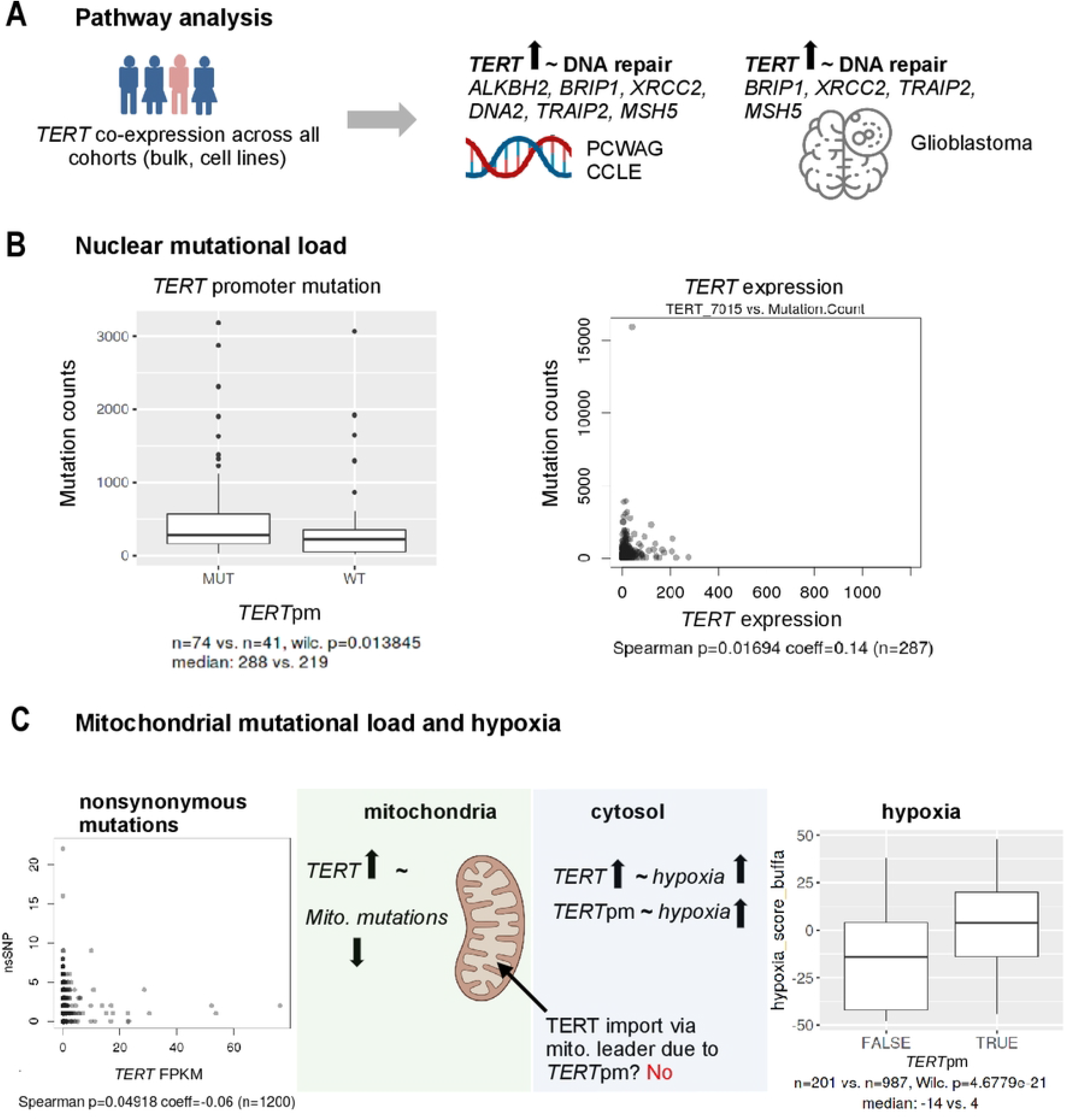
**A,** *TERT* expression and DNA repair genes associated across all cohorts. **B,** *TERT* alterations and exome mutation load in the SKCM cohort. **C,** *TERT* alterations, mitochondrial mutation load and hypoxia (PCAWG cohort). TPM: Transcripts per million; FPKM: Fragments per kilobase of transcript per million fragments mapped; nsSNP: Nonsynonymous mitochondrial SNP; *TERT*pm: *TERT* promoter mutations.

Moreover, in the SKCM cohort we did not find *TERT* promoter mutations associated with Breslow depth or age at diagnosis, and we did not find *TERT* expression associated with sex (SKCM, Liu), age at diagnosis (SKCM), or various clinicopathological variables of the Liu cohort, such as heterogeneity or total neoantigens (Supplementary Tables S3).

### *TERT* in melanoma subtypes

Because the BRAF-MAPK signaling pathway is frequently activated in melanoma [26], we analyzed *TERT* in melanoma subtypes with hotspot mutations in *BRAF, RAS (NRAS, KRAS, HRAS)* and *NF1* genes in the SKCM cohort. Melanomas with hotspot mutations carried *TERT* promoter mutations more often than triple wild-type melanomas (odds ratios >20, Supplementary Table S4B). With respect to *TERT* expression, we found significantly lower *TERT* expression in *BRAF* hotspot mutants than in *RAS* hotspot mutants (Wilcoxon p=0.0004, Supplementary Table S4A). The group of patients with *TERT* promoter mutations showed downregulated *TERT* expression in *BRAF* mutants and upregulated *TERT* expression in *RAS* mutants (mutated *RAS*: Wilcoxon p=0.024, mutated *BRAF:* Wilcoxon p=0.004, Supplementary Table S4C).

### Gene expression associated with *TERT* alterations

At first, we re-replicated the previously described association of higher *TERT* expression levels in the presence of *TERT* promoter mutations in three independent cohorts (Supplementary Table 8A). Since alternative splicing due to mutation of the *TERT* promoter could result in varied cellular functions of TERT beyond telomerase activity [17], we analyzed the expression of *TERT* isoforms in the presence of *TERT* promoter mutations. However, none of the isoforms were consistently regulated depending on the *TERT* promoter status (Supplementary Table S8A, Supplementary Fig. S8).

Next, we aimed to identify other cellular signaling pathways that may be driven by or co-regulated with either *TERT* promoter mutations (Supplementary Table S5) or *TERT* expression (Supplementary Table S6) followed by an enrichment analysis of associated gene expression.

Associations with the presence of *TERT* promoter mutations revealed an overlap of 30 genes in the three bulk RNA-seq cohorts (in melanoma and across entities), however from those, only six genes were consistently up- or down-regulated (up: *ARL17A, FBF1, KAZALD1, PRAF2*; down: *PLEKHB2, RASGRP3*; Supplementary Table S5). Only one gene (*PRAF2*) showed a consistent up-regulation also in the cell line data comprising other cancer entities (CCLE cohort). Enrichment analysis of the 30 genes associated with *TERT* promoter mutations showed overlaps for ‘cilium organization’; ‘plasma membrane bound cell projection assembly’ and ‘actin cytoskeleton’ (Supplementary Table S7A). To elucidate whether our findings are transferable to brain tumors, we extended our analyses to glioblastoma, in which *TERT* promoter mutations are highly recurrent [3] and are currently being integrated as biomarkers in diagnostic procedures [27]. Analysis of genes associated with *TERT* promoter mutations in 28 bulk tumor samples from the glioblastoma subcohort in PCAWG showed that of the six genes consistently associated with *TERT* promoter mutations in all bulk RNA-Seq cohorts, only *ARL17A* was upregulated in glioblastoma (Supplementary Table S5A).

In the context of *TERT* expression, 17 genes were consistently upregulated with *TERT* expression in all seven bulk RNA-seq cohorts, comprising melanoma and a cross-entity cancer cohort, as well as a cross-entity cell line cohort (Table 2, Supplementary Table S6). We confirmed that these *TERT* correlates were mostly tumor cell-specific genes as within melanoma, 15 genes were predominantly expressed in malignant tumor cells (Table 2, scRNA-seq). Enrichment analysis of the 17 genes upregulated with *TERT* expression revealed ‘DNA repair’, ‘double-strand break repair’, and ‘replication fork processing’ (Fig. 3A, Supplementary Table S7A). We replicated most of these associations in glioblastoma (Klughammer cohort) where 14 of 17 genes were also positively associated with *TERT* expression, including four genes related to DNA repair (*XRCC2, MSH5, TRAIP, BRIP1*, Supplementary Table S6).

**Table 2.** Genes repeatedly associated with TERT mRNA in 9 cohorts (Supplementary Table S6).

| Overlap of bulk RNA-seq and CCLE cohort | Overlap with malignant single-cell | Consistent up or downregulated | Function |
| --- | --- | --- | --- |
| 17 | 15 |  |  |
| FBXW9 | FBXW9 | Up | Substrate-recognition component of the SCF (SKP1-CUL1-F-box protein)-type E3 ubiquitin ligase complex |
| KIF14 | KIF14 | Up | Microtubule motor protein that binds to microtubules with high affinity through each tubulin heterodimer and has an ATPase activity (By similarity), plays a role in many processes like cell division, cytokinesis and also in cell proliferation and apoptosis |
| KLHL23 | KLHL23 | Up | Kelch-Like Family Member 23 (no function available) |
| MSH5 | MSH5 | Up | Involved in DNA mismatch repair and meiotic recombination processes |
| PUS7 | PUS7 | Up | Pseudouridylate synthase that catalyzes pseudouridylation of RNAs, acts as a regulator of protein synthesis in embryonic stem cells by mediating pseudouridylation of RNA fragments derived from tRNAs |
| SMARCA4 | SMARCA4 | Up | Involved in transcriptional activation and repression of selected genes by chromatin remodeling (alteration of DNA-nucleosome topology), component of SWI/SNF chromatin remodeling complexes |
| STIL | STIL | Up | Immediate-early gene, plays an important role in embryonic development as well as in cellular growth and proliferation, its long-term silencing affects cell survival and cell cycle distribution as well as decreases CDK1 activity correlated with reduced phosphorylation of CDK1 |
| TMEM201 | TMEM201 | Up | Involved in nuclear movement during fibroblast polarization and migration |
| TRAIP | TRAIP | Up | E3 ubiquitin ligase required to protect genome stability in response to replication stress |
| TTK | TTK | Up | Phosphorylates proteins on serine, threonine, and tyrosine, probably associated with cell proliferation |
| UCKL1 | UCKL1 | Up | May contribute to UTP accumulation needed for blast transformation and proliferation |
| XRCC2 | XRCC2 | Up | Involved in the homologous recombination repair (HRR) pathway of double-stranded |
| SMG5 | SMG5 | Up | Plays a role in nonsense-mediated mRNA decay |
| DNA2 | DNA2 | Up | Key enzyme involved in DNA replication and DNA repair in nucleus and mitochondria |
| ALKBH2 | ALKBH2 | Up | Dioxygenase that repairs alkylated DNA and RNA containing 1-methyladenine and 3-methylcytosine by oxidative demethylation |
| BRIP1 |  | Down | DNA-dependent ATPase and 5' to 3' DNA helicase required for the maintenance of chromosomal stability |
| ARGHGEF39 |  | Up | Rho Guanine Nucleotide Exchange Factor 3 |

*TERT* = telomerase reverse transcriptase; CCLE = cancer cell line encyclopedia
Note: Genes are sorted alphabetically and the first 12 genes were also found in glioblastoma
a Consistent up- or down-regulation: similar direction of association in all bulk, single-cell, and cell line cohorts

To further elucidate tumor cell-specific signaling pathways, we used single-cell RNA-seq data and analyzed genes co-expressed with *TERT.* Several cellular expression programs were repeatedly enriched, including the extracellular matrix, integrins, stromal cells, and smooth muscle cells (Supplementary Table S7B).

### Association of *TERT* and mitochondrial variables

To further elucidate the alternative role of TERT beyond telomere elongation in the molecular biology of mitochondria and reactive oxygen species (ROS) [15,28], we first tested associations of several ROS-related candidate genes with the presence of genetic alterations in *TERT*. However, we did not observe consistent upregulation or downregulation off the examined genes *NOX1-5, DUOX1/2, SOD1/2, OPA1, MFN1/2, PINK1, FUNDC1*, and *GPX4* in correlation with *TERT* promoter mutations or expression (Supplementary Table S8B,C). Yet, within our discovery approach, we identified a consistent upregulation of *DNA2* with higher *TERT* expression in several bulk tumor cohorts, potentially pointing to a role of *TERT* in DNA repair (Table 3).

We then analyzed the mutational load specifically in the mitochondria using the cross-entity PCAWG cohort (n=1200 with data on mitochondrial mutations, Supplementary Table S3C,E). Here, *TERT* expression correlated with fewer, especially nonsynonymous mitochondrial mutations (Wilcoxon p=0.007, Spearman p=0.049, coeff=−0.06, Supplementary Table 3D, Fig. 3C) in stark contrast to the increased overall, and mainly nuclear, mutations assessed above (in SKCM metastases, n=363 with data on overall mutations, Fig. 3B). *TERT* promoter mutation status was not associated with nonsynonymous or other mitochondrial mutations (Supplementary Table S3C, E).

We further tested whether the expression of the mitochondrial targeting sequence, which is required for the transport of TERT into the mitochondria [24], could be affected by *TERT* promoter mutations. Here, we did not detect a change in the proportion of expressed isoforms with mitochondrial targeting sequence (Supplementary Table 8A) and conclude that *TERT* promoter mutations do not promote or restrain transcripts with mitochondrial targeting (Fig. 3C).

### *TERT* alterations in hypoxia and chromothripsis

We next tested whether *TERT* promoter mutations and *TERT* expression correlated with hypoxia and chromothripsis in the PCAWG and SKCM cohorts (Supplementary Tables S3B-E). Ragnum hypoxia scores in melanoma patients (SKCM, Wilcoxon p=0.024, Supplementary Table S3C) and Buffa scores across entities (PCAWG, Wilcoxon p=4.678e-21, Fig. 3C) were higher in samples carrying a *TERT* promoter mutation. We also observed upregulation of *TERT* expression with all hypoxia scores in the melanoma cohort (Supplementary Table S3B) and with Buffa hypoxia scores across entities (Wilcoxon p=7.43e-22). We observed significantly more chromothripsis in samples carrying *TERT* promoter mutations and a higher median *TERT* expression in samples with chromothripsis (PCAWG, Supplementary Tables S3D-E).

## Discussion

### *TERT* and ICI therapy

*TERT* promoter mutations have been discussed as potential prognostic factors in metastatic melanoma [5,6,8,29,30] and as potentially predictive most recently under BRAF/MEK [7] and anti-CTLA4 therapy [9]. TMB is discussed as a biomarker for ICI success [31,32] and *TERT* promoter mutations showed association with higher TMB [9], however, multivariate approaches controlling for TMB were missing. Our analyses show that, when controlling for mutational load and prior MAPK inhibitor therapy, the association of *TERT* promoter mutations in the tested anti-CTLA4 and non-ICI cohorts weakened or was absent. Hence, we believe that the extent to which *TERT* promoter mutations govern survival under anti-CTLA4 therapy is not sufficiently clear [7,8]. In addition, since elevated *TERT* expression was not associated with survival, we conclude that melanoma differs from other cancer entities that showed an association between *TERT* levels and prognosis in lung cancer, oral squamous cell carcinoma, and colorectal carcinoma [10–12]. However, in these studies, TMB was not assessed, to our knowledge, and could possibly improve multivariate analyses. A striking correlation of *TERT* expression in our study was, however, a higher estimate of tumor-infiltrating CD4+ T cells, which were previously associated with improved clinical outcome of ICI therapy [33]. There is evidence for a baseline TERT-specific CD4+ T cell immune response, occurring in over 50% of melanoma patients, rising to 80% in ICI responders [34,35]. Consequently, a higher CD4+ T cell response, possibly explained by an anti-TERT T-helper type 1 response, could define a subgroup of patients with better ICI therapy response. However, our analyses of single-cell expression showed that, if malignant cells expressed higher levels of *TERT*, the corresponding CD4+ T cells from these tumors expressed exhaustion markers at higher levels (*CTLA4, PDCD1/PD1, HAVCR2/TIM3* and *TIGIT*, not *LAG3*, Fig. 1B). Therefore, in *TERT*-high hypoxic melanoma, dysfunctional CD4+ T cells may accumulate employing these immuno-suppressive checkpoints [36]. As previously suspected, this could argue for a defect in anti-TERT CD4+ response during melanoma progression [35] and may explain why in our analysis, patients with high CD4+ T cell estimates and increased *TERT* expression did not show a survival benefit.

### *TERT* expression in metastases from thin primaries

*TERT* promoter mutations were previously described to upregulate *TERT* expression [37,38] involving the formation of ETS/TCF transcription factor binding sites [2,37], and have been found in primary and metastatic melanoma. Here, metastatic melanomas showed higher *TERT* expression if they originated from thinner primaries, prompting the question of increased *TERT* expression in efficient early metastasis. This is especially relevant as *TERT* expression has been shown to be involved in most steps of the invasion-metastasis cascade [39]. The local invasion of primary melanoma cells might be regulated through increased *TERT* expression affecting cellular programs, such as NF-kB or metalloproteinases in addition to canonical telomerase activity [40,41]. As the gene expression enrichment of specifically malignant melanoma cells revealed that *TERT* expression is co-regulated with integrins, TGF-beta, and interleukins, a *TERT* expression shift may be involved in remodeling the extracellular matrix, which is seen as a driving force for cancer stemness [42] and would aid invasion. Yet it remains unclear whether *TERT* expression was elevated already in the thinner primary tumors corresponding to the metastases measured for RNA (Fig. 2B). Higher Breslow depth was previously shown to be associated with the presence of *TERT* mutations in primary tumors [5,29]. However, this association was not apparent in our analysis, probably because only thicker primaries were selected for the RNA-seq experiment in the SCKM cohort.

In conclusion, our data indicate that *TERT* expression may contribute to early metastasis from thin primaries, potentially through extracellular matrix remodeling, without a clear role for promoter mutations in this respect.

### Hotspot mutations, mutational load and DNA repair

Initially, we confirmed the known co-occurrence of *TERT* promoter mutations and hotspot mutations in *BRAF*, *RAS* and *NF1* as previously described for melanoma [5]. While, thus a co-mutation of the *TERT* promoter and members of the MAPK pathway is evident, *TERT* expression varied significantly between mutants, showing lower *TERT* expression in *BRAF* mutants compared to *RAS* mutants. Moreover, our finding that *BRAF* mutants exhibited lower *TERT* expression compared with non-*BRAF* mutants in *TERT* promoter-mutated samples somewhat contradicts the proposed model of *TERT* reactivation in *BRAF*-mutated samples through acquisition of *TERT* promoter mutations, subsequent binding of TCF, and thus increased *TERT* expression [43]. Since co-mutations were only available in one cohort, further investigations may improve our understanding of the MAPK pathway and *TERT* activation via *TERT* promoter mutations.

As a source of hotspot mutations, mutational load is also correlated with *TERT* expression in metastatic melanomas here and previously described[9]. The increased co-expression of DNA repair-related genes suggests that *TERT* expression is involved in establishing genomic stability under replication stress since *TERT* expression regulates the chromatin state and DNA damage responses in normal human fibroblasts [44]. Also beyond melanoma, co-expression of *TERT* and DNA repair genes was shared in various cancer entities including GBM, however, no common gene expression patterns were observed for *TERT* promoter mutations. To date, studies of *TERT* alterations in GBM have focused on the fairly frequent promoter mutations. Hence, our results may spark interest in assessing *TERT* expression levels in GBM to test sensitivity to DNA damage [45]. This could be relevant in the context of chemo- and radiotherapy independent of *TERT* promoter mutations, since *TERT* expression is also mediated via rearrangements and hypermethylation [46].

Moreover, our data support the view that this non-canonical function of *TERT* could be especially relevant under hypoxic conditions and after chromothripsis, where *TERT* expression was increased. In line with the proposed concept of a two-step activation of *TERT* [37], the up-regulation of *TERT* expression in later stages of melanoma development could aid the survival of melanoma cells in hypoxic tumors and metastasis to other sites.

### *TERT* and mitochondrial stability

In cancer cells exposed to hypoxia, ROS levels produced by mitochondria are increased[47]. Moreover, *TERT* has previously been shown to be involved in the protection of cells from ROS-mediated oxidative stress, delaying aging processes supposedly via TERT activity in mitochondria [15]. As these effects were described predominantly in non-cancer tissues, we aimed to identify potential mitochondrial correlates of *TERT* expression in cancer and, specifically, in melanoma. While at first, we did not find associations of *TERT* with a set of fifteen ROS-related candidate genes, our less-biased discovery approach identified a consistently upregulated *DNA2*, known to be involved in DNA repair in the nucleus and mitochondria [48]. This is in line with *TERT* translocation into mitochondria to protect mitochondrial DNA from damage and implicates a dedicated repair gene in the described process [15,24]. The correlation between *TERT* expression and fewer nonsynonymous mitochondrial mutations further supports this rationale. It contrasts though, with the fact that *TERT* expression and *TERT* promoter mutations were associated with more overall mutations in the nuclear genome. Therefore, DNA repair via *TERT* and possibly *DNA2* may be mitochondria-specific. Interestingly, differing mutational processes have already been described in colorectal and gastric cancers, where microsatellite instability was observed in the nuclear, but not in the mitochondrial genome [49]. Thus, it would be highly interesting to explore the cellular localization of TERT and DNA2 in various cancers and in the context of hypoxia.

In conclusion, our analyses did not support a significant association between *TERT* genetic alterations and survival in melanoma patients. Hence, its presumed role as a biomarker, especially for anti-CTLA4 or MAPK inhibitor therapy, needs further validation. In addition, our data provide evidence for dysfunctional CD4+ T cells in TERT-highly hypoxic melanomas as a possible reason for the lack of survival benefit. Since melanoma metastases with high *TERT* expression were shown to have originated from thinner primary tumors, we conclude that upregulation of *TERT* expression may contribute to metastasis. This is also supported by the involvement of *TERT* expression in the dynamics of the extracellular matrix. We add support to the non-canonical function of *TERT* expression related to mitochondrial and nuclear DNA stability even across cancer entities, especially under hypoxic conditions and after chromothripsis. In summary, our results demonstrate the possible role of *TERT* expression in metastasis formation and invasion in melanoma and highlight its role in DNA repair mechanisms across cancer entities.

## Financial support

This work was funded by the Deutsche Forschungsgemeinschaft (DFG, German Research Foundation, HO 6389/2-2, SCHA 422/17-2, ‘KFO 337’ - 405344257).

## Author Contributions

Conceptualization: Christina Kuhn, Susanne Horn Resources: Sven-Holger Puppel, Susanne Horn Data curation: Christina Kuhn, Sven-Holger Puppel, Susanne Horn Software: Susanne Horn

Formal analysis: Christina Kuhn, Susanne Horn

Supervision: Susanne Horn, Torsten Schöneberg

Funding acquisition: Susanne Horn, Dirk Schadendorf, Torsten Schöneberg

Validation: Christina Kuhn, Susanne Horn

Investigation: Christina Kuhn, Susanne Horn Visualization: Christina Kuhn, Susanne Horn Methodology: Christina Kuhn, Susanne Horn

Writing original draft: Christina Kuhn, Susanne Horn, Jaroslawna Meister, Sophia Kreft, Mathias Stiller, Anne Zaremba, Björn Scheffler, Vivien Ullrich

Project administration: Susanne Horn, Dirk Schadendorf, Torsten Schöneberg

Writing-review and editing: all authors

## Acknowledgements

We thank Udo Stenzel for help with the data analysis.

## Supplementary tables and figures

**Supplementary Table S1.** Multivariate analysis using Cox proportional hazard models for progression free survival (PFS) (Supplementary Table S1A) and overall survival (OS) (Supplementary Table S1B) subsetted for ICI therapy type in cohorts Liu, Gide, Hugo, Riaz, Van Allen and SKCM.

**Supplementary Table S2.** Significant immune signatures in SKCM associated with *TERT* expression and comparison against other bulk RNA-seq cohorts (Gide, Hugo, Liu, Riaz, Van Allen).

**Supplementary Table S3.** Wilcoxon tests for group comparisons and Spearman correlations of clinical variables against *TERT* expression and *TERT* promoter mutation in Liu, SKCM and PCAWG cohorts (Supplementary Table S3A-E).

**Supplementary Table S4.** Mutation subtypes against *TERT* expression (Supplementary Table S4A) and *TERT* promoter mutation status (Supplementary Table S4B) in the SKCM melanoma cohort. Effect of hotspot mutations on *TERT* expression in *TERT* promoter mutated samples (Supplementary Table S4C).

**Supplementary Table S5.** Genes associated with *TERT* promoter mutation status for the Van Allen, SKCM, PCAWG, and CCLE cohorts, and overlap of cohorts (Supplementary Table S5A-E).

**Supplementary Table S6.** Genes associated with *TERT* expression in each cohort (Gide, Riaz, Van Allen, Hugo, Liu, SKCM, PCAWG, CCLE, single-cell, and Klughammer) and overlap of cohorts (Supplementary Table S6A-K).

**Supplementary Table S7.** Enrichment analysis of 17 genes associated with *TERT* expression and 30 genes associated with *TERT* promoter mutation in bulk tumors and cell lines (Supplementary Table S7A), and for the top 100 genes associated with *TERT* expression in single-cells (Supplementary Table S7B).

**Supplementary Table S8.** Influence of *TERT* promoter mutation on *TERT* isoforms, mitochondrial targeting sequence (Supplementary Table 8A), associations of ROS-related genes with *TERT* expression levels (Supplementary Table 8B), and *TERT* promoter mutation status (Supplementary Table 8C) in Gide, Riaz, Hugo, Liu, Van Allen, SKCM, PCAWG, CCLE cohorts, and single-cell.

**Supplementary Figure S1.** Overview of the analysis workflow.

**Supplementary Figure S2.** Survival analysis under anti-CTLA4 ICI with ipilimumab in the melanoma VanAllen cohort and the non-ICI cohort (SKCM). Comparison of *TERT* low vs. high patients and *TERT* promoter mutation status.

**Supplementary Figure S3.** Survival analysis under anti-PD1 ICI with nivolumab in three melanoma cohorts (Liu, Gide, Riaz). Comparison of *TERT* low vs. high patients.

**Supplementary Figure S4** Survival analysis under anti-PD1 ICI with pembrolizumab in three melanoma cohorts (Liu, Gide, Hugo). Comparison of *TERT* low vs. high patients

**Supplementary Figure S5.** Univariate and multivariate survival analysis of reanalyzed melanoma MSK Impact 2017 subcohort. Comparison of *TERT* promoter mutated patients vs. non mutated.

**Supplementary Figure S6.** Survival of selected patients with a high CD4+ Th1 xCell signature comparing *TERT* low vs. high patients in melanoma cohorts (SKCM, Van Allen, Riaz, Gide, Hugo, Liu).

**Supplementary Figure S7.** *TERT* isoform expression in normal control tissues (Gtex database).

**Supplementary Figure S8.** *TERT* isoform with increased expression in the presence of promoter mutation.

**Supplementary Figure S9.** Breslow depth with *TERT* expression and *TERT* promoter mutations including outliers.

## Notes

**Conflict of interest statement**, The authors declare no potential conflicts of interest.

### Competing Interest Statement

The authors have declared no competing interest.

